# “Conformational dynamics of C1r inhibitor proteins from Lyme disease and relapsing fever spirochetes”

**DOI:** 10.1101/2023.03.01.530473

**Authors:** Sourav Roy, Charles E. Booth, Alexandra D. Powell-Pierce, Anna M. Schulz, Jon T. Skare, Brandon L. Garcia

## Abstract

Borrelial pathogens are vector-borne etiological agents of Lyme disease, relapsing fever, and *Borrelia miyamotoi* disease. These spirochetes each encode several surface-localized lipoproteins that bind to components of the human complement system. BBK32 is an example of a borrelial lipoprotein that protects the Lyme disease spirochete from complement-mediated attack. The complement inhibitory activity of BBK32 arises from an alpha helical C-terminal domain that interacts directly with the initiating protease of the classical pathway, C1r. *Borrelia miyamotoi* spirochetes encode BBK32 orthologs termed FbpA and FbpB, and these proteins also inhibit C1r, albeit via distinct recognition mechanisms. The C1r-inhibitory activities of a third ortholog termed FbpC, which is found exclusively in relapsing fever spirochetes, remains unknown. Here we report the crystal structure of the C-terminal domain of *B. hermsii* FbpC to a limiting resolution of 1.5 Å. Surface plasmon resonance studies and assays of complement function demonstrate that FbpC retains potent BBK32-like anti-complement activities. Based on the structure of FbpC, we hypothesized that conformational dynamics of the complement inhibitory domains of borrelial C1r inhibitors may differ. To test this, we utilized the crystal structures of the C-terminal domains of BBK32, FbpA, FbpB, and FbpC to carry out 1 µs molecular dynamics simulations, which revealed borrelial C1r inhibitors adopt energetically favored open and closed states defined by two functionally critical regions. This study advances our understanding of how protein dynamics contribute to the function of bacterial immune evasion proteins and reveals a surprising plasticity in the structures of borrelial C1r inhibitors.

## Introduction

Pathogenic bacteria must evade the immune system of their hosts in order to establish infection. To this end, many bacterial pathogens have evolved mechanisms to disrupt the powerful bacteriolytic activity and immune surveillance capacity of the complement system (1–8). For example, the etiologic agents of borreliosis are master manipulators of the complement cascade due to the production of numerous anti-complement proteins on the bacterial surface (3, 9, 10). As such, the causative agent of Lyme disease, *Borreliella (Borrelia) burgdorferi*, has become a powerful model system for understanding mechanisms of microbial complement evasion (3, 9, 11, 12). Although less studied, several complement-targeting lipoproteins have also been described for other agents of borreliosis including tick-borne relapsing fever spirochetes (*i.e.*, *Borrelia hermsii, Borrelia turicatae, etc.*) (13–17), louse-borne relapsing fever spirochetes (*i.e., Borrelia recurrentis*) (18, 19) and *Borrelia miyamotoi* spirochetes (BMD) (20–23).

An example of a borrelial surface exposed lipoprotein that mediates complement inhibition is BBK32 (24–27). BBK32 is produced *in vivo* and is important for optimal infection in a murine model of Lyme borreliosis (28, 29). BBK32 is a multifunctional protein that interacts with host glycosaminoglycans and fibronectin using separate intrinsically disordered N-terminal binding sites (24, 26, 28, 29). BBK32 also blocks the classical pathway (CP) of the complement system by specifically binding and inhibiting the initiating protease C1r via its carboxy terminal alpha helical domain (*i.e.*, BBK32-C) (3, 30–32). BBK32-C forms a high-affinity interaction with the serine protease domain of human C1r (C1r-SP) and is capable of binding both zymogen and activated forms of C1r (30, 31). Small angle X-ray scattering (SAXS) and X-ray crystallography structures of the complex between fragments of human C1r and BBK32-C, coupled with extensive site-directed mutagenesis studies, further defined the protein-protein interface formed between BBK32-C and C1r-SP (32). These studies revealed that BBK32 occludes the S1 subsite of the C1r active site with a cluster of residues located near a small loop that connects BBK32 alpha helix 1 and alpha helix 2 (32). At this BBK32/C1r-SP binding site – which we will refer to as the “K1 site” here forward – a key arginine residue of BBK32 (*i.e.*, R248) was shown to be critical to both C1r-binding affinity and complement inhibitory activity (32).

A second BBK32/C1r-SP binding surface, which we term here the “K2 site”, involves BBK32 residues on alpha helix 5, typified by lysine 327 of BBK32 (K327). The precise inhibitory functions of the K2 site are less clear. However, the K2 site, which is ∼25 Å away from the K1 site, interfaces with non-active site residues on a surface loop of C1r known as the B-loop. The importance of the K2 site is accentuated by the BBK32-K327A mutant, which exhibits an ∼150-fold decrease in C1r binding affinity and significantly reduced complement inhibitory activity (32). Furthermore, natural mutations for another residue of the K2 site (*i.e.*, BBK32-E324) were identified in a homolog from avian-associated *B. garinii* (*i.e.*, BGD19) that had significantly reduced activity against human complement (31). Nonetheless, in BBK32, only simultaneous mutation of both R248 and K327 within the K1 and K2 site respectively (*i.e.,* BBK32-R248A-K327A) resulted in a protein that lacked measurable C1r binding and caused a complete loss of serum resistance in a serum-sensitive *B. burgdorferi* background (32). Thus, the K1 and K2 sites each contribute significantly to the activity of BBK32 against human complement.

Relapsing fever spirochetes encode BBK32-like proteins that organize phylogenetically into three families known as FbpA, FbpB, and FbpC (20, 33). Whereas Lyme disease spirochetes encode a single BBK32 protein, relapsing fever and *B. miyamotoi* spirochetes encode either two, or in some cases, all three Fbp proteins (20, 33). The C1r-binding properties and inhibitory mechanisms of Fbp proteins have been characterized for FbpA and FbpB from *B. miyamotoi* (20). FbpA interacts with C1r and inhibits the CP in a similar manner to BBK32, including retaining the ability to bind both zymogen and active forms of C1r (20). Mutations to the homologous K1 and K2 site residues in FbpA (*i.e.*, FbpA-R264A-K343A) lost all C1r-binding and complement inhibitory activity, which provided independent validation of the importance of these conserved sites. Interestingly, FbpB bound only to the active form of C1r, and the crystal structure of the C-terminal region of FbpB revealed significant deviations in the secondary structure of the FbpB K2 site relative to both BBK32 and FbpA (20). FbpB also lacks the conserved Lys residue in the K2 site that is shared between BBK32 and FbpA (*i.e.*, BBK32-K327/FbpA-K343) and is instead replaced with a small insertion loop that interrupts the 5^th^ alpha helix. Furthermore, several residues on this K2 site loop in FbpB are apparently flexible, as evidenced by a lack of identifiable electron density in crystallographic experiments utilizing this domain of FbpB (20).

In contrast to BBK32, FbpA, and FbpB, little is known about the C1r-inhibitory properties of FbpC proteins. However, FbpC proteins from *B. recurrentis* and *B. duttonii* have been reported to be involved in complement evasion by specific binding of the endogenous complement regulators C1 esterase inhibitor (C1-INH) and C4b-binding protein (C4BP) (18). This protein was designated CihC and a putative binding site for C1-INH and C4BP was mapped to residues 145 to 185 (18). In a follow up study, CihC/FbpC orthologues were studied in *B. hermsii*, *B. parkeri,* and *B. turicatae* (34). While none of these proteins retained the C1-INH or C4BP-binding properties of *B. recurrentis* CihC/FbpC, four out of the five proteins bound specifically to fibronectin (34). Finally, *B. hermsii* FbpC from strain HS1 is known to be expressed during a murine model of infection (33). This protein, initially referred to as BHA007, was shown to bind to fibronectin and C4BP, and the authors proposed the BBK32/Fbp nomenclature which we have retained here (33). Collectively, these reports show that a subset of FbpC proteins use N-terminal binding sites to recruit host complement regulators and serve as a fibronectin adhesin (18, 33, 34). However, none of these prior studies addressed the function of the C-terminal domain of FbpC.

In this study, we report the structural and functional characterization of the C-terminal region of a representative FbpC family member from *B. hermsii,* termed FbpC-C hereafter. Our biochemical, microbiological, and biophysical studies demonstrate that *B. hermsii* FbpC is a potent inhibitor of the classical pathway of human complement. Surprisingly, however, the high-resolution crystal structure of FbpC-C exhibited a striking difference in the conformation of the K1 site compared to BBK32, FbpA, and FbpB. Based on this observation, we tested the hypothesis that protein dynamics play a previously unrecognized role in the structure and function of borrelial C1r inhibitors by carrying out detailed analysis of 1 µs MD simulations and biochemical and biophysical assays utilizing site-directed mutants of FbpC-C.

## Results

### The crystal structure of the C-terminal domain of B. hermsii FbpC

The crystal structures of the C-terminal complement inhibitory domains of *B. burgdorferi* BBK32-C (PDB ID:6N1L), *B. miyamotoi* FbpA-C (PDB ID:7RPR), and *B. miyamotoi* FbpB-C (PDB ID:7RPS) have all been previously reported (20, 31). However, a representative structure of the C-terminal region of an FbpC family member remained elusive. We overcame this by identifying a crystallizable construct of *B. hermsii* strain HS1 FbpC corresponding to residues 212-374 (*i.e.*, FbpC-C) that diffracted to 1.50 Å limiting resolution (**Fig. 1A, Fig. S1A, Table 1**). Protein crystals of FbpC-C grew in space group P 2_1_ with a single molecule in the asymmetric unit. Initial phases were obtained by molecular replacement using a model derived from AlphaFold2 (35). Iterative rounds of refinement resulted in a final model (*R_work_* = 17.7% and *R_free_* = 19.8%, **Table 1**) that was deposited into the Protein Data Bank (PDB ID: 8EC3).

**Figure 1.**
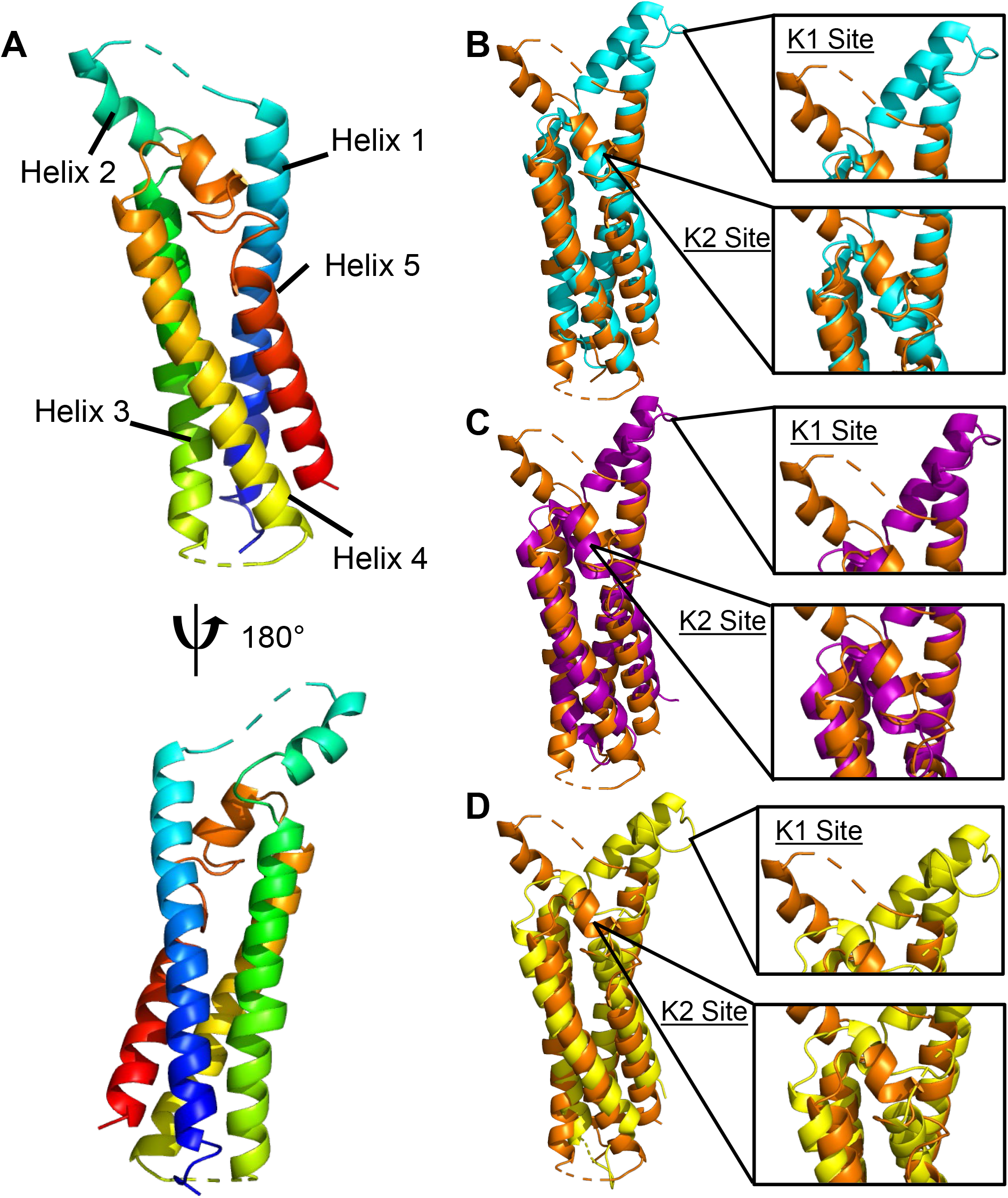
*B. hermsii* FbpC-C crystal structure. **(A)** The crystal structure of *B. hermsii* FbpC-C_212-374_ (from strain HS1) at a limiting resolution of 1.50 Å (PDB ID: 8EC3). The model is colored as a spectrum with the N-terminus in blue and C-terminus in red. Structural overlays of FbpC-C (orange) with **(B)** BBK32-C (cyan, PDB ID: 61NL), **(C)** FbpA-C (purple, PDB ID: 7RPR), and **(D)** FbpB-C (yellow, PDB ID: 7RPS). The K1 and K2 sites are defined as the areas of BBK32-C that interact with the S1 pocket and B-loop of C1r respectively (32).

**Table 1.** Data collection and refinement statistics.

|  |  |
| --- | --- |
|  | FbpC (212-374) |
| **Data collection** |  |
| Space group | P 1 21 1 |
| Cell dimensions |  |
| a, b, c, Å | 38.24, 46.56, 45.19 |
| $\alpha, \beta, \gamma, ^\circ$ | 90.00, 110.80, 90.00 |
| Resolution, Å | 31.28-1.50 (1.55-1.50) |
| $R_{\text{pim}}$ | 0.032 (0.218) |
| $R_{\text{meas}}$ | 0.087 (0.509) |
| $CC_{1/2}$ | 0.989 (0.870) |
| I/σI | 19.4 (2.0) |
| Completeness, % | 98.7 (89.2) |
| Redundancy | 7.0 (4.2) |
| **Refinement** |  |
| Resolution, Å | 28.35-1.50 (1.55-1.50) |
| No. reflections | 22,687 (1,796) |
| $R_{\text{work}}/R_{\text{free}}$ | 0.1770/0.1975 |
| No. non-hydrogen atoms | 1,393 |
| Protein | 1,235 |
| Water | 157 |
| B-factors |  |
| Protein | 30.38 |
| Water | 39.40 |
| Rmsd |  |
| Bond lengths, Å | 0.010 |
| Bond angles, ° | 1.05 |

*Values in parentheses refer to the highest-resolution shell.*

### Structural similarity between FbpC-C and BBK32-C/FbpA-C/FbpB-C

Analysis of the FbpC-C structure revealed that it shares a common four-helix bundle fold with BBK32-C, FbpA-C, and FbpB-C (**Fig. 1**) (20, 31). Our previous structural and biochemical studies identified two regions of BBK32-C that are critical for interaction with human C1r (31, 32). The first, denoted here as the “K1 site”, is found on a small loop that connects alpha helix 1 and alpha helix 2, and in BBK32 harbors a key arginine residue (*i.e.*, R248). A homologous arginine residue (*i.e.*, R264) was also shown to be important for C1r-binding by FbpA-C, and FbpB-C presents R309 in a structurally similar position to both BBK32-C and FbpA-C (20). Intriguingly, a sequence alignment to BBK32-C, FbpA-C, and FbpB-C predict that a histidine residue in FbpC-C (*i.e.*, H252) takes the place of the key K1 site arginine residues noted above for BBK32-C/FbpA-C/FbpB-C.

Surprisingly, the crystal structure of FbpC-C deviated from BBK32-C, FbpA-C, and FbpB-C in the conformation of K1 site in two major ways (**Fig. 1**). First, the relative angle of alpha helices 1-2-3 is ∼75° in FbpC-C, whereas this angle is ∼20-25° in BBK32-C, FbpA-C, and FbpB-C. The open conformation of the FbpC-C helix was not predicted in the AlphaFold2 model, where it instead adopts a closed conformation (∼23°) (**Fig. S1B**). While the AlphaFold2 model was characterized by overall very high confidence, as judged by predicted local distance difference test (pLDDT) ≥ 90 (35), we noted that both the K1 and K2 sites showed reduced pLDDT values, dipping near or below the low confidence threshold (*i.e.*, pLDDT ≤ 70) (**Fig. S1C**). The second notable feature of this region of the FbpC-C crystal structure was the lack of electron density corresponding to the residues that connect the first and second alpha helix (residues M_247_-RMSGHST_254_) (**Figs. 1A, S1A**). Flexible regions of proteins are often not visualized in crystal structures (36), and thus, this result suggests that FbpC-C harbors a longer flexible loop connecting the first and second alpha helix that was not found in the crystal structures of other BBK32/Fbp family members (20, 31).

A second site was shown to be critical for BBK32-C interaction with C1r and inhibition of complement, termed the “K2 site”, which involves surface residues presented near a kink in the C-terminal 5^th^ alpha helix (32). In BBK32, the K2 site includes a key lysine residue that interacts with the B-loop of C1r (*i.e.*, K327) (32). A homologous lysine residue in FbpA-C (*i.e.*, K343) is also important for C1r-binding and inhibition (20). In contrast, a homologous lysine residue is absent from FbpB-C and is instead replaced with a loop structure (*i.e.,* Q394 to Q405) that interrupts the 5^th^ alpha helix (20). In the FbpC-C crystal structure the K2 site exhibits an unwinding of 5^th^ alpha helix into a larger loop structure (residues 344-354), more similar to what was observed for this site in FbpB-C (20). However, unlike FbpB-C, and despite the noted differences in secondary structure, FbpC-C presents a surface exposed lysine residue at a structurally homologous position to BBK32 K327 and FbpA K343 (**Fig. 1B-D**) (20, 31). Collectively, this analysis reveals that, while the core folds are similar, distinct structural features are concentrated at the K1 and K2 sites across the family of borrelial C1r inhibitors, and in particular for *B. hermsii* FbpC.

### B. hermsii FbpC binds both active and zymogen forms of C1r and inhibits human complement

Site-directed mutants and naturally occurring sequence variations at the K1 and K2 sites of BBK32 and Fbps alter their C1r-binding and complement inhibitor activities (20, 30, 31). Considering this, and given the structural differences noted above, we next tested the C1r-binding properties of FbpC-C. Previously, we have shown that BBK32-C and FbpA-C bind with high affinity to both zymogen and active forms of human C1r, while FbpB-C is selective for the activated form (20, 30–32). To determine if FbpC-C binds zymogen and/or active human C1r we generated a biosensor by immobilizing FbpC-C on a surface plasmon resonance (SPR) sensorchip and analyzed its binding to purified human zymogen C1r or activated C1r (**Figs. 2A,B**) (20). Like BBK32-C and FbpA-C, FbpC-C interacted with both zymogen and active forms of purified human C1r with high affinity (*K*_D_ values of 1.2 and 0.3 nM, respectively) (**Figs. 2A, B**).

**Figure 2.**
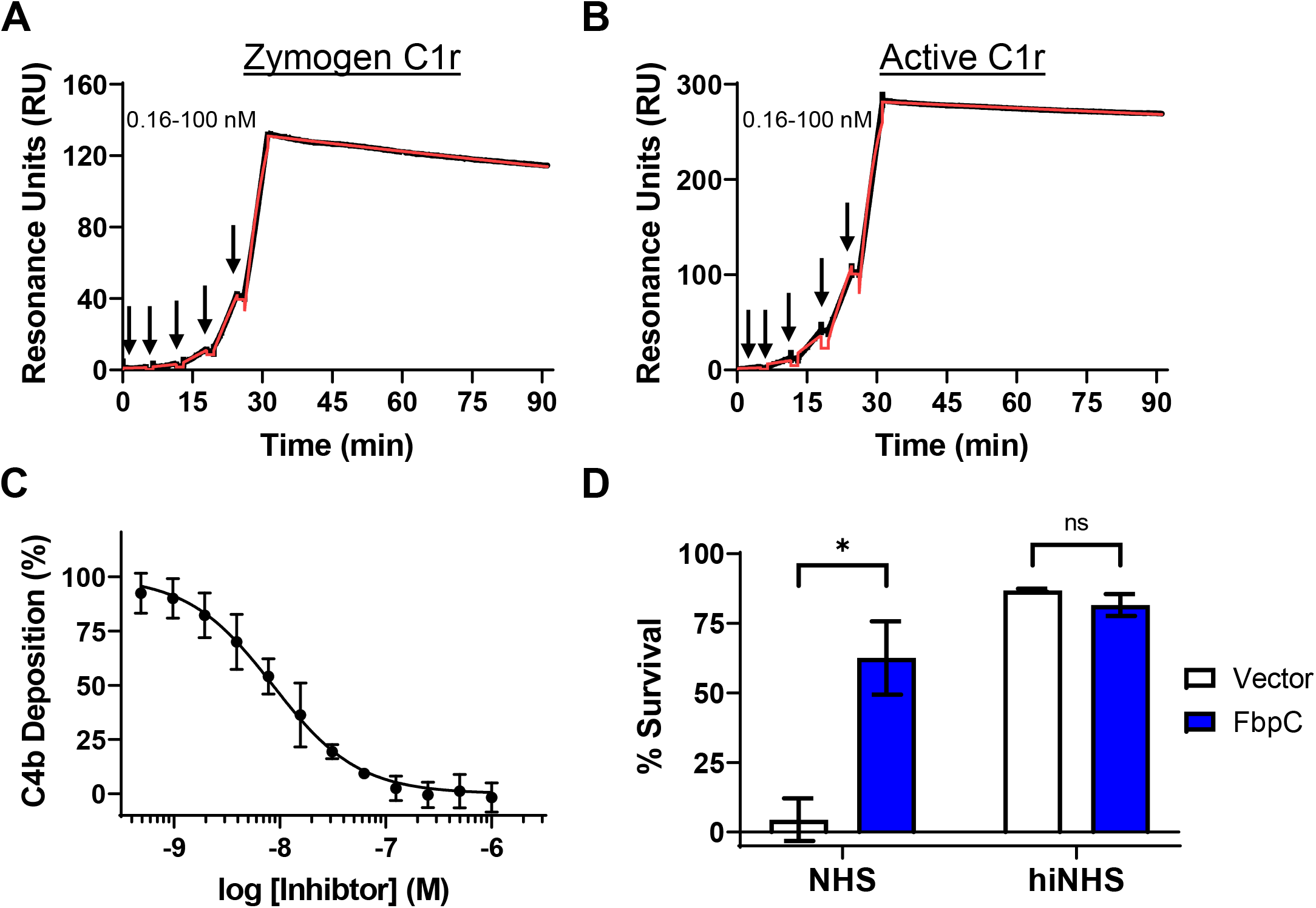
*B. hermsii* FbpC C1r-binding and complement activity assays. Single-cycle SPR binding assays using a 0.16-100 nM fivefold dilution series of **(A)** zymogen or **(B)** active C1r over FbpC-C. Sensorgrams are a representative injection series from three replicates with the raw sensorgrams drawn in black and kinetic fits traced in red. Calculated *K*_D_ values and association (*k*_a_) and dissociation (*k*_d_) rate constants for each analyte are shown in **Table 2**. **(C)** FbpC-C dose-dependently inhibits C4b deposition in a CP-specific complement ELISA. ELISAs were performed in triplicate followed by a non-linear variable slope regression fit to determine IC_50_ values present in **Table 2**. **(D)** Serum sensitivity assay demonstrating that *B. burgdorferi* B314 ectopically expressing *B. hermsii fbpC* significantly protects against complement-mediated bacteriolysis when compared to a vector-only strain. Heat inactivated normal human serum (hiNHS) was used a control. Statistical significance was determined using a two-way ANOVA and defined as p<0.05.

**Table 2.** C1r binding and inhibitory CP ELISA data.

*Values in parentheses refer to the highest-resolution shell.*
| **Table 2. C1r binding and inhibitory CP ELISA data.** |  |  |  |
| --- | --- | --- | --- |
|  | FbpC-C | FbpC-C R248A | FbpC-C H252A |
| SPR: Zymogen C1r $K_D$ (nM) | $1.2 \pm 0.034$ | $13 \pm 6.6$ | N.D. |
| $k_a$ (1/Ms) | $4.4 \times 10^4 \pm 1.2 \times 10^4$ | $6.9 \times 10^4 \pm 7.0 \times 10^4$ | N.D. |
| $k_d$ (1/s) | $5.4 \times 10^{-5} \pm 1.4 \times 10^{-5}$ | $6.3 \times 10^{-4} \pm 3.6 \times 10^{-4}$ | N.D. |
| SPR: Active C1r $K_D$ (nM) | $0.3 \pm 0.014$ | $0.19 \pm 0.048$ | $2.4 \pm 0.12$ |
| $k_a$ (1/Ms) | $4.8 \times 10^4 \pm 0.048 \times 10^4$ | $4.5 \times 10^4 \pm 0.52 \times 10^4$ | $5.1 \times 10^4 \pm 0.63 \times 10^4$ |
| $k_d$ (1/s) | $1.4 \times 10^{-5} \pm 0.076 \times 10^{-5}$ | $9.4 \times 10^{-6} \pm 3.1 \times 10^{-6}$ | $1.2 \times 10^{-4} \pm 0.19 \times 10^{-4}$ |
| CP ELISA $IC_{50}$ (nM) | 8.6 (7.5-9.8) | 23 (20-26) | 300 (240-400) |

N.D. stands for not determined.

Next, we determined if FbpC-C inhibits the progression of the human CP. Consistent with the observed high affinity binding to C1r (**Fig. 2**), FbpC-C inhibited the human CP in a dose-dependent manner with an IC_50_ of 8.6 nM (**Fig. 2C**). We then asked if ectopic expression of full-length FbpC on a surrogate borrelial strain could prevent complement-mediated bacteriolysis. To this end, we used a high-passage *B. burgdorferi* strain (*B. burgdorferi* B314) that lacks all linear plasmids and is highly sensitive to human serum-based complement-mediated killing (20, 32, 37). *B. burgdorferi* B314 producing full-length, surface-expressed FbpC **(Fig. S2)** showed a significant increase in resistance to human serum killing when compared to a *B. burgdorferi* B314 vector-only control (**Fig. 2D**), indicating that FbpC is sufficient to protect this strain from complement-mediated bacteriolysis. FbpC-C also was unable to inhibit the lectin pathway of complement at concentrations up to 1 μM, indicating that FbpC-C is CP-specific like BBK32-C (**Fig. S3**) (30). Taken together, these results show that *B. hermsii* FbpC binds to human C1r, inhibits the CP, and protects *B. burgdorferi* from complement-mediated killing.

### MD simulations of borrelial C1r inhibitor proteins

With crystal structures of a representative family member of each borrelial C1r inhibitor in hand (PDB: 6N1L, 7RPR, 7RPS, 8EC3), we next set out to utilize molecular dynamics (MD) simulations to test the hypothesis that BBK32 and/or Fbps are structurally dynamic at either the K1 site, K2 site, or both functional sites. This hypothesis was primarily formed from the crystallographic studies of FbpC-C (lack of K1 site electron density) and FbpB-C (lack of K2 site electron density) (20). MD simulations predict how atoms in a protein move over time (38) and have become a powerful approach for understanding protein motions that occur on the ns-µs timescale, which include conformational changes of side chains and loops (38–41). To study these motions in borrelial C1r inhibitors, we carried out 1 µs MD simulations on *B. burgdorferi* BBK32-C, *B. miyamotoi* FbpA-C and FbpB-C, and *B. hermsii* FbpC-C in triplicate. Representative simulations for each inhibitor are shown in **Supplemental Movies 1-4**.

### Initial analysis of MD simulations

Root mean square deviation (RMSD) describes the average displacement of atoms at a snapshot in time relative to a reference frame (*i.e.*, the crystal structure) and is a measure of the stability of a given simulation (42). Thus, RMSD reports on global conformational changes of protein structure throughout the simulation. RMSD values of the alpha carbons (Cα RMSD) of BBK32-C, FbpA-C, FbpB-C and FbpC-C showed that most simulations converged after 100-200 ns, with FbpC-C simulations converging near 500 ns (**Figs. 3A-D**). BBK32-C demonstrated the lowest RMSD values, ranging across the three simulations at 1.70 – 2.52 Å, followed by FbpA-C (2.36 – 3.04 Å) and FbpB-C (2.25 – 2.60 Å). FbpC-C RMSD values were relatively higher over the other three inhibitors, ranging between 3.29 and 3.81 Å, suggesting an increase in overall structural flexibility of FbpC-C compared to BBK32-C, FbpA-C and FbpB-C.

**Figure 3.**
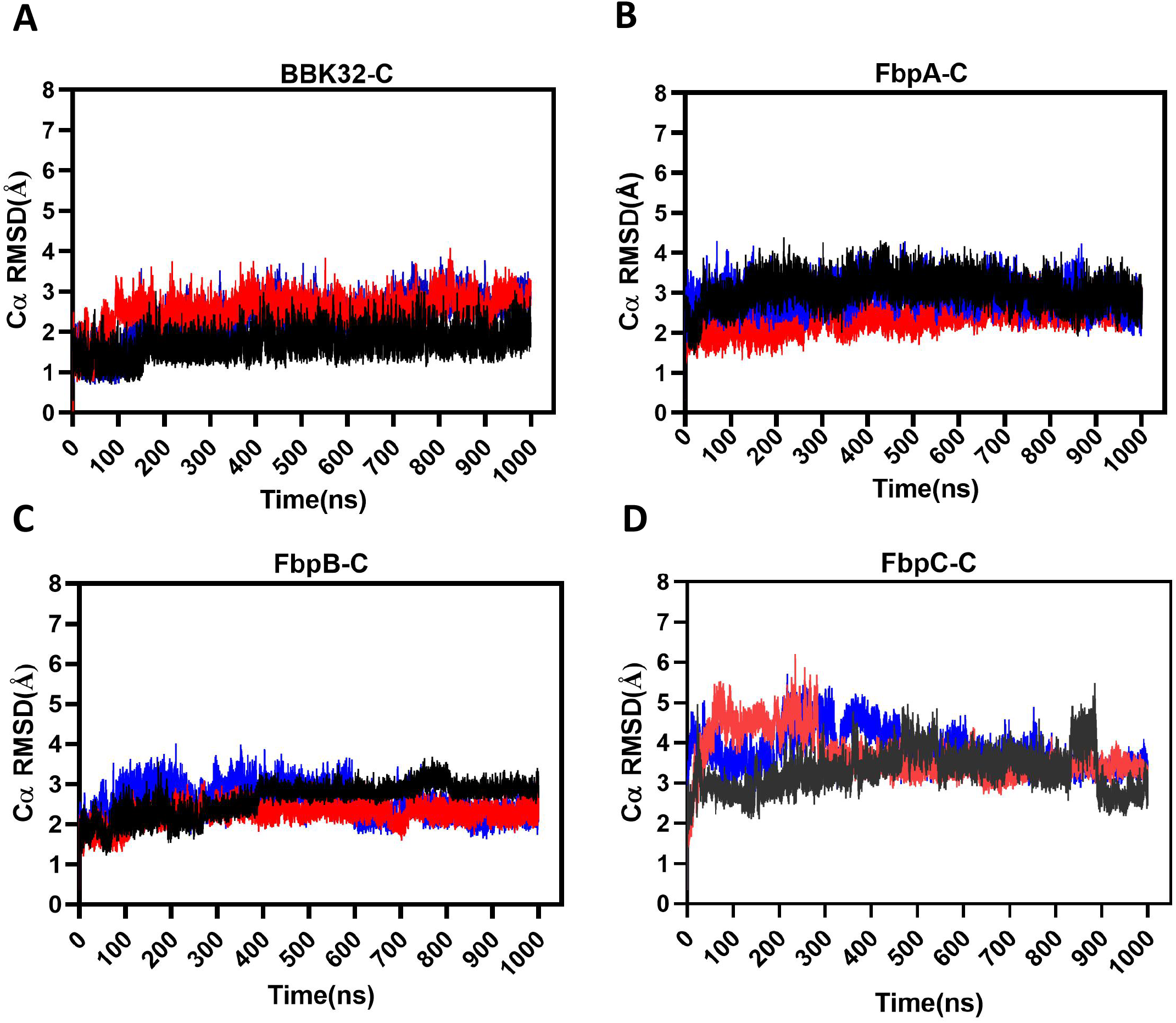
C*α* RMSD plots. The Cα root mean square deviation (RMSD) was tracked over the course of each 1 µs simulation for **(A)** BBK32-C, **(B)** FbpA-C, **(C)** FbpB-C and **(D)** FbpC-C. Simulation replicate 1 (black), 2 (blue) and 3 (red) are shown.

Next, we evaluated local residue flexibility by calculating root mean square fluctuations (RMSF). RMSF is conventionally calculated for groups of atoms that comprise a residue and are compared to the initial starting structure (43). Residue RMSF measures can identify amino acids that contribute most to the molecular motions of a protein in the timeframe of the simulation. As judged by residue RMSF values (**Fig. 4**), the K1 site residues form one of the most dynamic regions within each of the borrelial C1r inhibitor proteins. Interestingly, the K2 site is relatively inflexible in BBK32-C, FbpA-C, and FbpC-C, whereas this site has increased dynamics in FbpB-C. In all simulations, common areas of high residue RMSF values are found at the N- and C-terminal residues of each protein, and on the loop that connects alpha helices 3 and 4 (**Fig. 4**). Taken together, analysis of 1 µs MD simulations of BBK32-C, FbpA-C, FbpB-C, and FbpC-C suggests that a major functional site (*i.e.*, the K1 site) is dynamic in all four inhibitors with the most flexibility observed in FbpC-C. While relatively less flexible in all proteins compared to the K1 site, the K2 site in FbpB-C is more dynamic when compared to BBK32-C, FbpA-C, and FbpC-C, which is consistent with the lack of electron density for this loop in the prior FbpB-C crystal structure (20).

**Figure 4.**
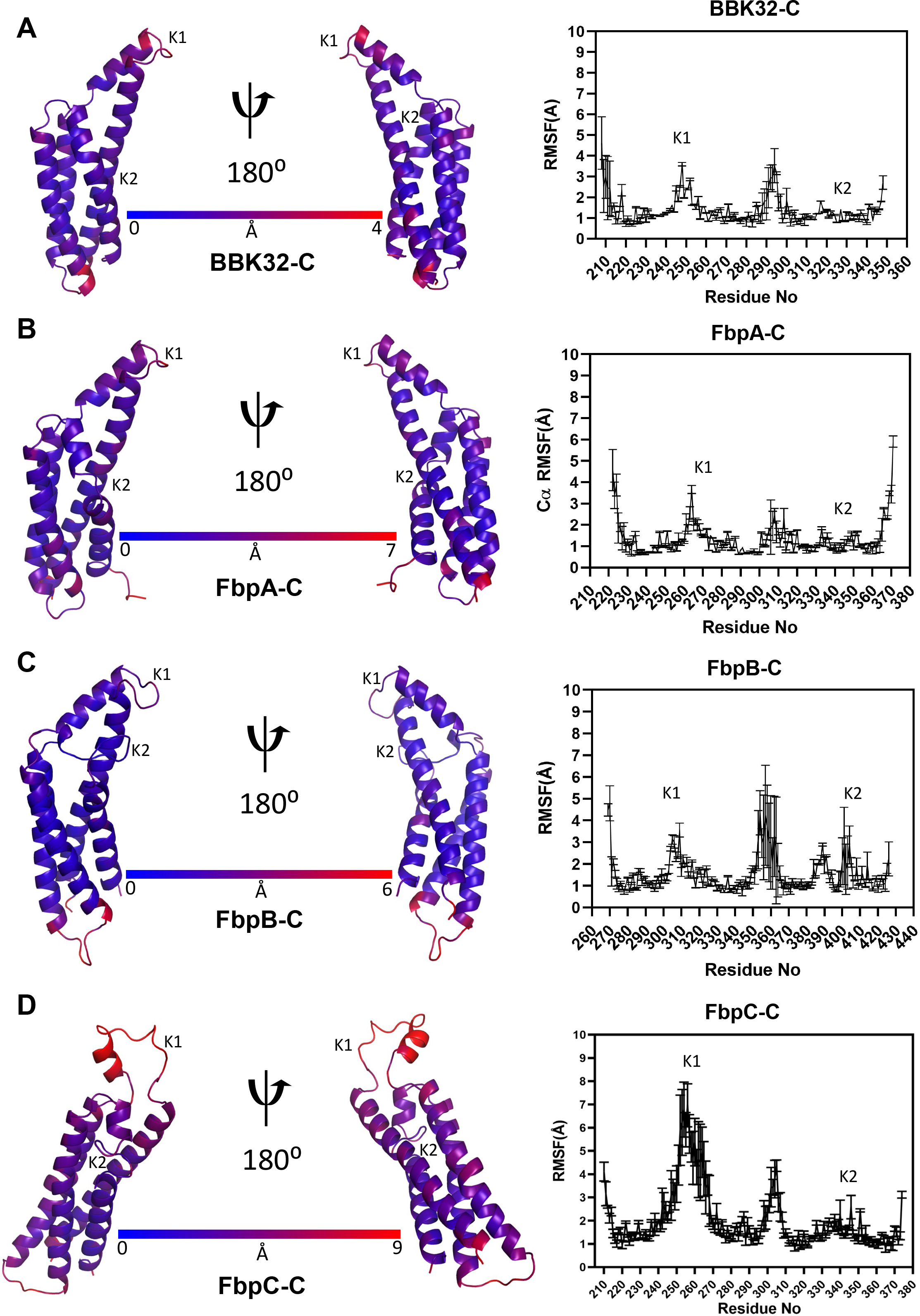
RMSF measurements. Residue root mean square fluctuation (RMSF) values from a representative MD simulation of **(A)** BBK32-C, **(B)** FbpA-C, **(C)** FbpB-C and **(D)** FbpC-C were mapped onto structural models of each protein. Dark blue indicates the least flexible residues while dark red indicates the most flexible residues within each protein. On the right panels, RMSF values are shown as an average and standard deviation of three replicate simulations. The location of the K1 and K2 sites are marked.

### K1 and K2 sites of borrelial C1r inhibitors display anti-correlated motions

To further investigate the dynamics of the K1 and K2 sites in borrelial C1r inhibitors, we carried out all atom normal mode analysis (**Fig. S4**) (44, 45). Normal mode analysis has been used to detect larger scale motions in proteins which are involved in biological functions (46–52). Separate all atom normal modes for BBK32-C, FbpA-C, FbpB-C and FbpC-C showed that the largest amplitude motions localize to the K1 site with other small amplitude normal modes distributed throughout each of the proteins. Interestingly, normal modes at the K2 sites in all proteins are significantly less in amplitude but are directed opposite to that of the corresponding K1 site (**Fig. S4**).

To further investigate the relative conformations of the K1 and K2 sites in these proteins, we carried out dynamic cross correlation matrix (DCCM) analysis of the MD simulations. DCCM measures positional displacements of the Cα atoms at a sample rate of 100 ps, which provides a graphical representation of time-correlated motions of residues within a given protein (53–56). Correlated motions (**Fig. 5**, red) are apparent for the helices that form the core of each borrelial C1r inhibitor. Strikingly, in each protein the residues that comprise the K1 and K2 sites (yellow boxes) move in an anticorrelated manner relative to one another (**Fig. 5**, blue). This analysis is consistent with the results of the normal mode analysis (**Fig. S4**), and, taken together, suggests that borrelial C1r inhibitor proteins may utilize a highly flexible K1 site and anti-correlated motion at the K2 site to generate variation in the distance between the two functional sites.

**Figure 5.**
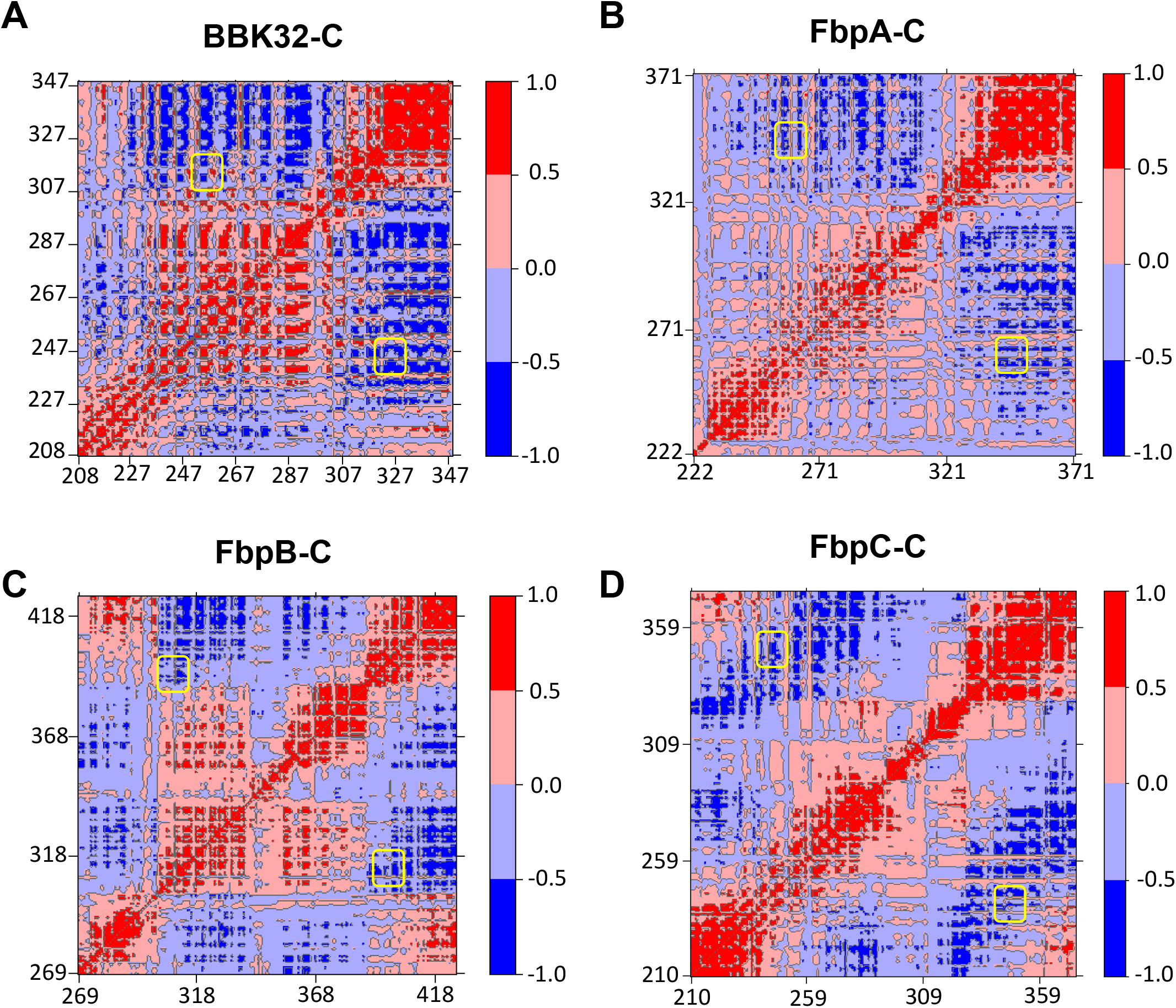
C*α* DCCM Maps for borrelial C1r inhibitors. Dynamic cross-correlation matrices (DCCM) maps are shown for representative simulations of **(A)** BBK32-C, **(B)** FbpA-C, **(C)** FbpB-C and **(D)** FbpC-C using a relative scale. Positive values (red) represent correlated motions, while negative values (blue) represent anti-correlated motions. Regions of the maps corresponding to K1 and K2 sites are highlighted by yellow boxes.

### Conformational plasticity of the K1 and K2 sites in borrelial C1r inhibitors

Analysis of the MD simulations led to a new hypothesis that the two major C1r-binding surfaces on free borrelial C1r inhibitors (*i.e.*, K1 and K2 sites) – which are separated in space by ∼25-30 Å – are sampling a larger range of conformations in their unbound forms, particularly for FbpC-C. To test this, we first defined the distance between the K1 and K2 sites in each protein by measuring the distance between two representative residues from each site using the Cα atom (*i.e.*, main chain distance) or a distal atom from the side chain as follows: i) BBK32-C; K1: R248, K2: K327; ii) FbpA-C; K1: R264, K2: K343; and iii) FbpB-C; K1: R309, K2: N402. As noted above, the FbpC sequence differs in this area from the other three proteins and a sequence alignment predicts that H252 is analogous to the K1 site arginine residues from BBK32-C, FbpA-C, and FbpB-C. However, we noted that an arginine residue is also present on the flexible K1 site loop in FbpC-C (*i.e.*, M_247_-<u>R</u>MSG<u>H</u>ST_254_; note that H252 is also underlined). Thus, we tracked two measurements for FbpC-C: 1) K1: R248; K2: K345 and 2) K1: H252; K2: K345.

Using these measures, we first analyzed how K1-K2 site distances change during the simulation (**Fig. S5).** Distances between K1 and K2 sites in BBK32-C ranged from 22-30 Å (main chain; average = 25.3 Å) and 12-35 Å (side chain; average = 23.3 Å). We then projected two-dimensional free energy surfaces using main chain and side chain K1-K2 site distances as order parameters (57, 58). This approach was taken to assess the relative K1-K2 site distances of energetically favorable conformations within unbound forms of each protein. This analysis suggests that BBK32-C samples energetically favorable conformations that place the K1 and K2 sites between main chain distances of 24-27 Å and side chain distances of 18-25 Å apart (**Fig. 6A, dark blue**). Similar results were obtained for FbpA-C with the most favorable conformations leading to K1-K2 distances between 25-27 Å (main chain) and 20-28 Å (side chain) (**Fig. 6B**). FbpB-C produces a similar shape of the free energy surface plot and indicates that the most favorable K1-K2 site distances range from 19-22 Å (main chain) and 22-26 Å (side chain) (**Fig 6B-C**). These results reveal that in the unbound form, BBK32-C, FbpA-C, and FbpB-C sample a similar range of energetically favorable states that modulate the relative positions of the two major C1r-binding surfaces, which presumably adopt a fixed distance in the bound state.

**Figure 6.**
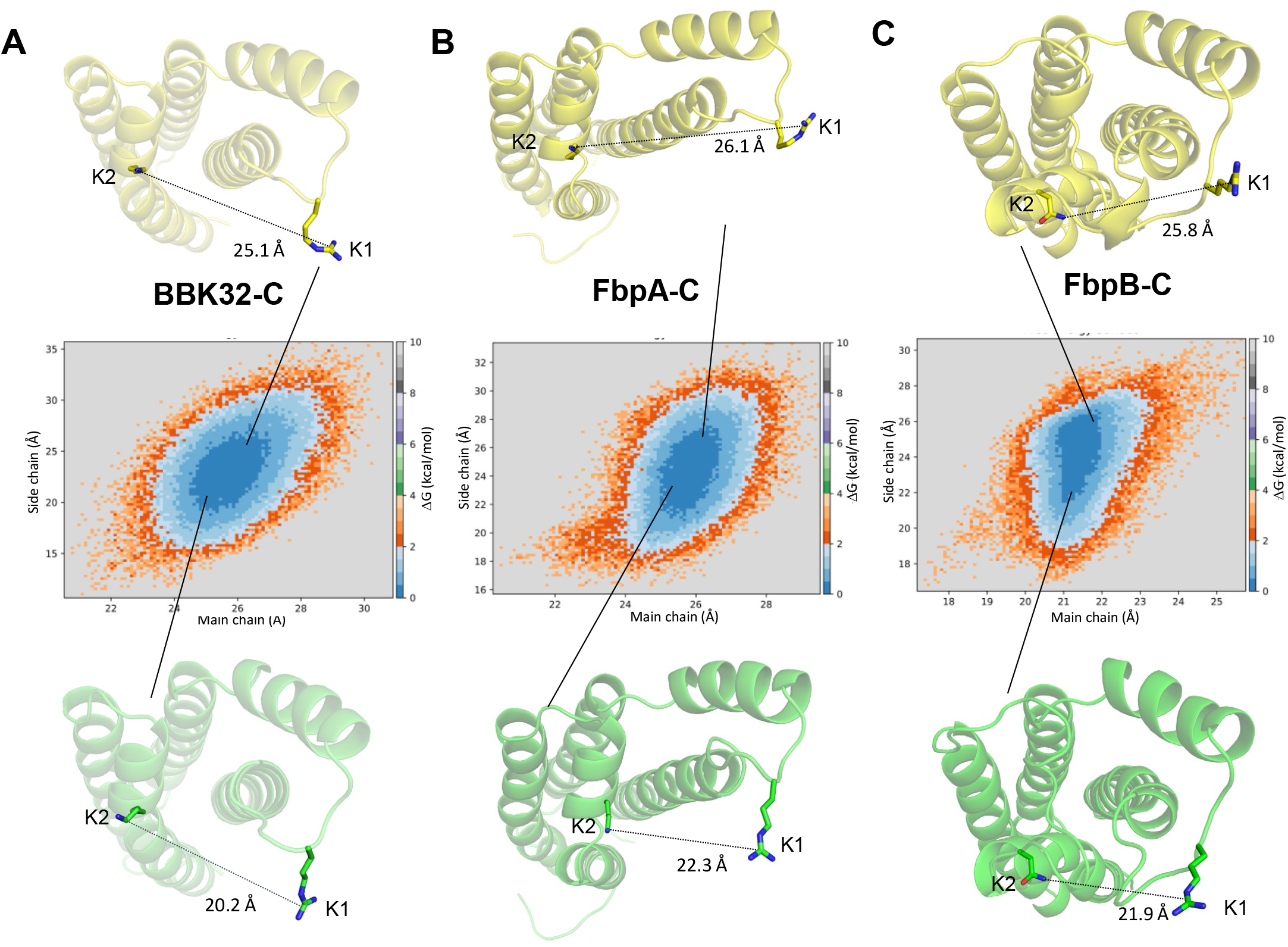
Free energy surfaces of borrelial C1r inhibitors in relationship to the main chain and side chain distances between the K1 and K2 sites. Two-dimensional free energy surfaces are shown for (A) BBK32-C, (B) FbpA-C, and (C) FbpB-C. Distance between the K1 and K2 sites at main chain and side chain atoms are used as order parameters. The residues used for these measurements are defined as BBK32-C (K1: R248, K2: K327); FbpA-C (K1: R264, K2: K343); FbpB-C (K1: R309, K2: N402). Side chain distances are defined as distances between the K1 site Arg-CZ atom and K2 site Lys-NZ atom in BBK32-C and FbpA-C. In FbpB-C, side chain distances were defined as distances between K1 Arg-CZ and ND2 atom of N402. A relative energy scale is shown on the right y-axis with dark blue indicating the lowest energy conformations. Representative conformations from the edge of each energy minima are shown with K1 and K2 side chain distance measurements drawn.

Unlike the other energy contours, FbpC-C exhibits three clear energy minima when R248 is used in the measurement of the K1-K2 site distance (**Fig. 7A**). The largest basin, represented by the cyan structure in **Fig. 7A**, involves energetically favorable conformations that produce K1-K2 site distances of 18-20 Å (main chain) and 18-23 Å (side chain). A more closed conformation (represented by the green structure) and more open conformation (represented by the yellow structure) are also apparent (**Fig. 7A**). A distinct free energy surface is also seen when H252 is used to measure the K1-K2 site distance in FbpC-C (**Fig. 7B**). Two energy minima are observed that collectively span 21-30 Å (main chain) and 14-25 Å (side chain). This suggests that FbpC-C adopts energetically favorable conformations that produce a wide range of distances of the K1 and K2 sites. Also, in contrast to the other three inhibitors, the secondary structure of the K1 site undergoes considerable rearrangement during the time course of the simulation. As an example, the conformation of the more closed state (**Fig. 7, green**) involves a complete unwinding of alpha helix 2. Likewise, the helical structure of the K1 site observed in the more open state (**Fig. 7, yellow**) differs from that found in the central energy basin (**Fig. 7, cyan**).

**Figure 7.**
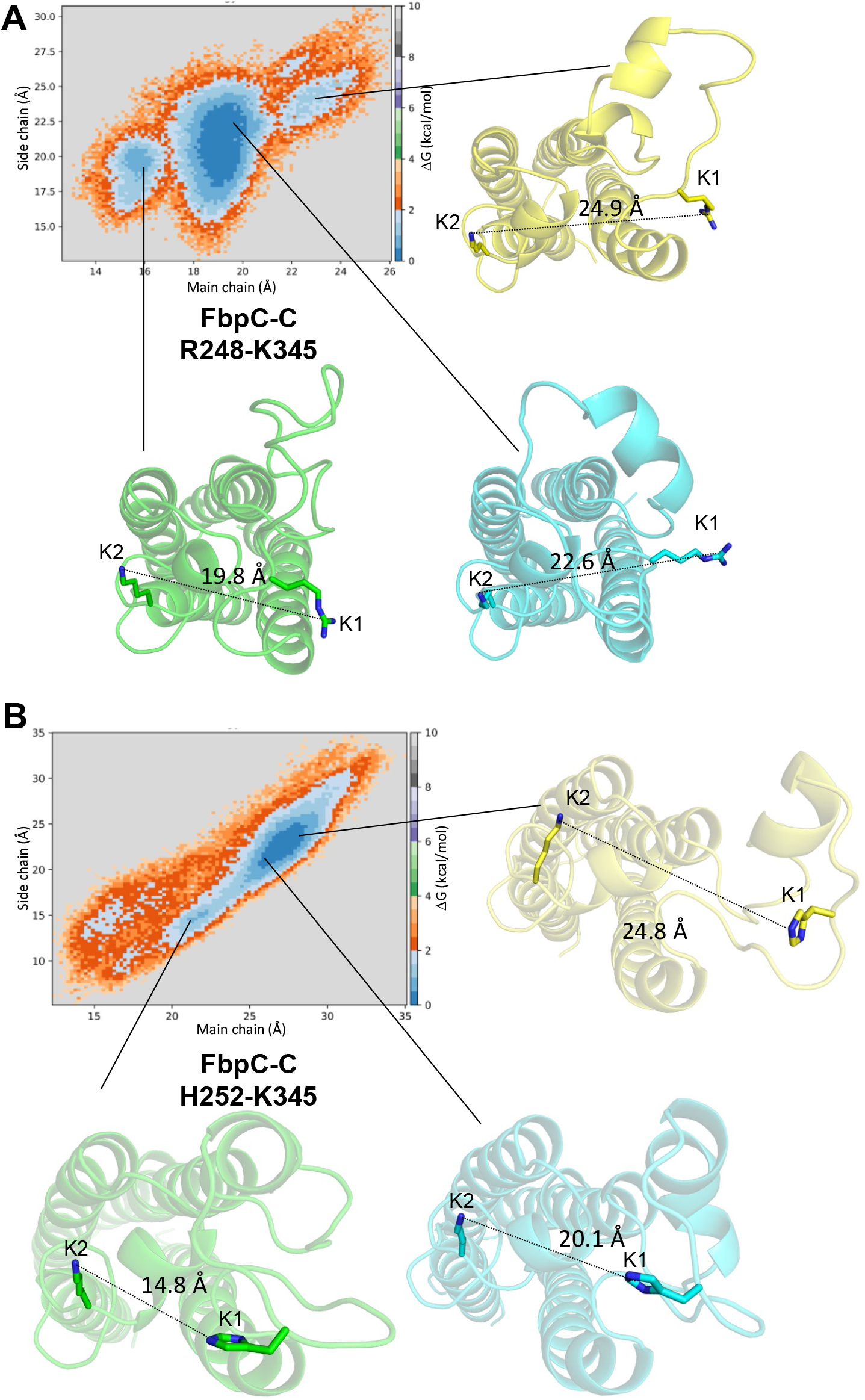
Free energy surfaces of FbpC-C. **A)** Two-dimensional free energy surfaces were generated for FbpC-C. Two sets of K1 and K2 main chain and side chain distances are used as reaction coordinates for generating such surfaces. **A)** R248 and K345 with side chain distances defined using the Arg-CZ atom of R248 and the NZ atom of K345. **B)** Free energy surfaces generated for FbpC-C using H252 using between NE2 atom of H252 to measure the side chain distance. Representative conformations are shown for three energy minima for the most closed (green), most open (yellow), and median distances (cyan).

### Dynamic K1 site residues are important for FbpC-C function

There was ambiguity in determining which residue within the dynamic K1 site of FbpC-C (i.e.: R248 or H252) was involved in C1r interactions based on sequence and structural alignments. Furthermore, the analysis conducted in **Figs. 7** and **S5** suggest that both residues can adopt conformations that would feasibly allow them to interact with C1r, as judged by the previous studies with BBK32, FbpA, and FbpB (20, 32). To test if R248 and/or H252 contribute to C1r-binding affinity we generated two site-directed alanine mutants, FbpC-R248A and FbpC-H252A, and assayed binding to C1r using SPR and the CP ELISA (**Fig. 8**). FbpC-R248A bound zymogen and active C1r with respective *K*_D_’s of 13 and 0.19 nM and dose-dependently inhibited the CP with an IC_50_ of 23 nM, an overall 2-3-fold difference in inhibition. In contrast, FbpC-C H252A exhibited a weaker affinity for active C1r (*K*_D_ = 2.4 nM) than wild-type FbpC (*K*_D_ = 0.3 nM) and even more so for zymogen C1r where a *K*_D_ could not be reliably determined (**Figs. 8A-B**). Consistent with this C1r-binding deficit, CP-specific ELISAs showed that the H252A mutation had a pronounced effect on CP inhibition with an IC_50_ value of 300 nM, compared to 8.6 nM for wild-type FbpC-C (**Fig. 2C** and replotted in **Fig. 8C** for comparison). These data highlight that both residues affect binding and inhibition of C1r to various degrees and underscore the importance that dynamic residues play for overall complement inhibitory properties of FbpC-C.

**Figure 8.**
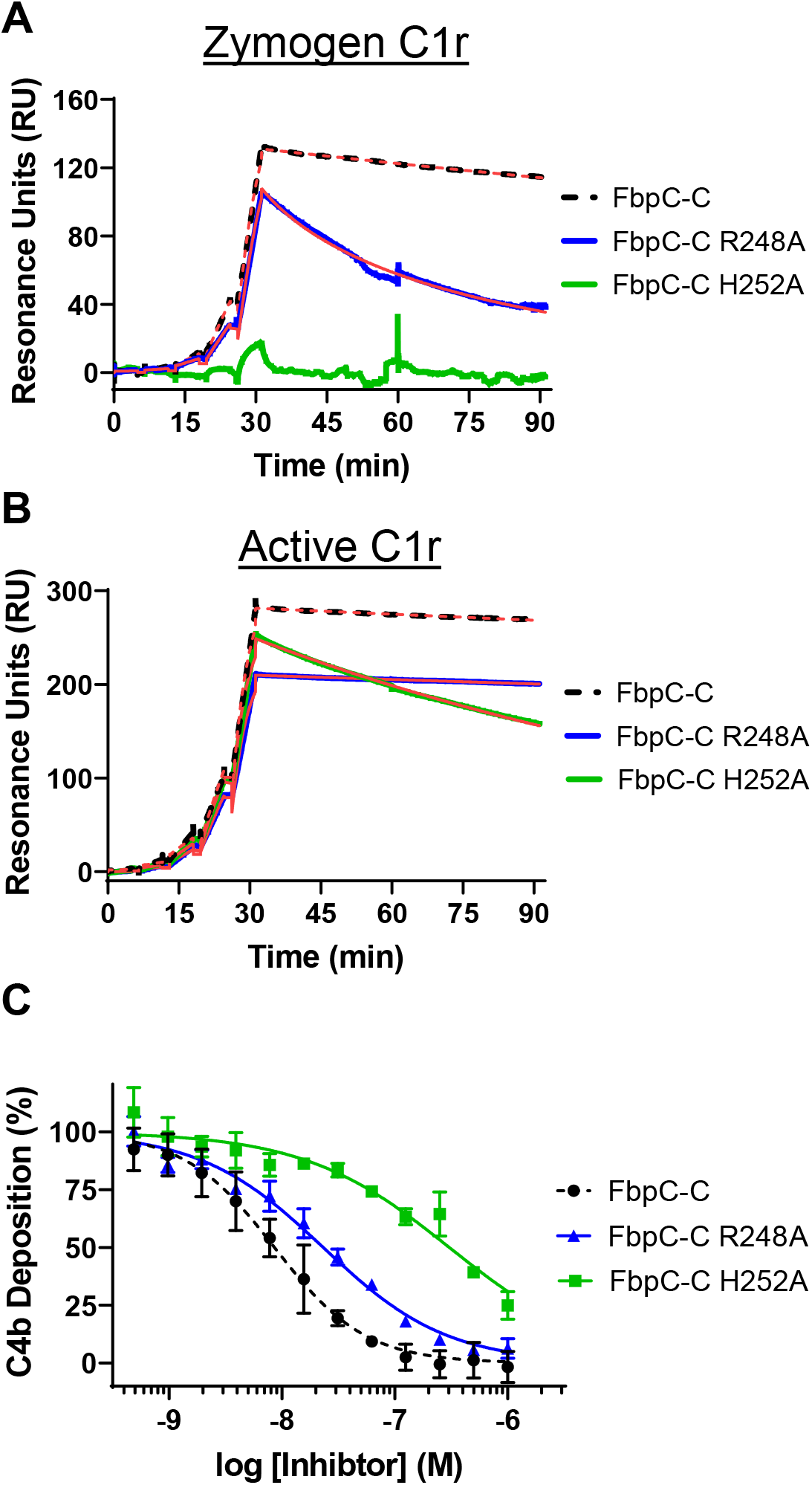
Assessing the function of the K1 site residues of FbpC-C. Single-cycle SPR binding assays using a 0.16-100 nM fivefold dilution series of **(A)** zymogen or **(B)** active C1r over FbpC-C R248A and H252A. Sensorgrams are a representative injection series from three replicates with the raw sensorgrams overlaid with kinetic fits traced in red. A kinetic fit for zymogen C1r binding to FbpC-C H252A was unable to be determined and is shown only as the raw sensorgram. Calculated *K*_D_ values and association (*k*_a_) and dissociation (*k*_d_) rate constants for each analyte were determined and are shown as the mean ± SD (**Table 2**). Representative sensorgrams for analogous experiments using “wild-type” FbpC-C are replotted from Fig. 2 for sake of comparison and are delineated by dashed lines. **C)** C4b deposition was dose-dependently inhibited in a CP-specific complement ELISA incubated with FbpC-C R248A and FbpC-C H252A. CP ELISAs were performed in triplicate followed by a non-linear variable slope regression fit to determine IC_50_ values (**Table 2**). Representative dose-response inhibition curves for analogous experiments using “wild-type” FbpC-C are replotted from Fig. 2 for sake of comparison and are delineated by dashed lines.

## Discussion

To colonize and persist following infection, many bacterial pathogens produce outer surface-associated proteins that specifically interact with host proteins. These bacteria-host protein-protein interactions have diverse functions, including facilitating pathogen adherence to tissues, promoting invasion of cells, and providing protection against host immune defenses (1–4, 8, 17, 22, 59–61). Lyme disease spirochetes produce at least 80 outer surface lipoproteins, many of which are upregulated upon transmission from the vector to the host (62). While the functions of most of these proteins remain unknown, nearly a dozen have been shown to bind directly to protein components of the innate immune system known as the complement cascade (3, 9). Some of these complement evasion lipoproteins act as highly specific inhibitors of complement proteases, including a family of C1r inhibitors prototyped by *B. burgdorferi* BBK32 and the orthologous Fbp proteins of relapsing fever and *B. miyamotoi* spirochetes (18, 20, 30–33). In this study we begin to assess how sequence diversity across C1r borrelial inhibitors influences their structure and function.

Our previous investigation into the molecular basis for C1r inhibition by BBK32, FbpA, and FbpB, defined two C1r-interacting surfaces located on the alpha helical C-terminal region of each protein referred to herein as K1 and K2 sites. However, the structure and C1r inhibitory activity of a representative FbpC protein had not been determined. Interestingly, the crystal structure of *B. hermsii* FbpC-C, presented here, exhibited structural deviations at both the K1 and K2 sites compared to BBK32-C, FbpA-C, and FbpB-C (**Fig. 1**). In particular, the conformation of the K1 site involved a repositioning of alpha helix 2 coupled with the absence of strong electron density for eight residues that connect alpha helix 1 and alpha helix 2 which suggested this region was highly flexible. (**Fig. 1A**). We cannot rule out the influence of the crystallization process/conditions in stabilizing the conformation observed for the FbpC-C K1 site, but we note that BBK32-C, FbpA-C, and FbpB-C were all crystallized such that the K1 sites were in a more closed conformation. Despite the structural differences for FbpC-C, we showed that it retains high affinity for human C1r and inhibits human complement in a manner similar to BBK32 and *B. miyamotoi* FbpA (**Fig. 2**).

Our structure-based knowledge of bacteria-host protein-protein interactions is often derived from static three-dimensional models produced by X-ray crystallography. Powerful new machine learning-based approaches, such as AlphaFold2, also produce static representations of protein structures. However, proteins are dynamic entities, and their native conformational variations can dramatically influence their functions. Crystallographic B-factors – or the noted correlation of pLDDT values to protein order in AlphaFold2 models (63) – provides some information on protein motions. However, their utility is somewhat limited as protein dynamics occur across different timescales and are associated with different distances. Here we studied protein dynamics of C1r borrelial inhibitors using MD simulations, which allows for visualization of protein dynamics on the ns-µs timescale (38, 40). This enabled us to observe motions ranging from side chain fluctuations to larger conformational changes associated with loop movements and secondary structure changes, particularly at the K1 site (**Figs. 3-4**) (39, 40). Previous studies using MD have discovered novel functional domains of proteins (64) and aided in understanding how surface receptors transmit their signals within the cell (65). Regarding bacterial proteins, MD simulations have been used to reveal how polymorphisms in FnBPA of *Staphylococcus aureus* increases affinity for human fibronectin resulting in an increased ability to colonize medical implant devices (66). To our knowledge, the application of MD for borrelial proteins has been limited to studying DNA-protein interactions or periplasmic proteins (67, 68), and is thus a novel approach for the study of borrelial outer surface lipoproteins.

The crystal structure of FbpC-C (**Fig. 1**), and results from RMSF analysis (**Fig. 4**) for the MD simulations, suggested that the K1 sites of borrelial C1r inhibitors were highly dynamic. Interestingly, DCCM and normal mode analyses also indicated that the K1 and K2 site of each protein moved in an anti-correlated fashion (**Figs. 5, S4)**. Free energy surface analysis showed that BBK32 and Fbp proteins adopt energetically favorable “open” and “closed” states defined by the variation in distance between the functional K1 and K2 sites (**Fig. 6-7**). FbpC-C was particularly dynamic at the K1 site and our investigations into the flexible loop of the K1 site of FbpC-C by site-directed mutagenesis showed that the H252A had an approximately 35-fold decrease in CP inhibition and decreased binding to C1r (**Fig. 8**). In contrast, the R248A mutant protein exhibited an approximate three-fold decrease in inhibition of the CP and was decreased in its ability to bind zymogen C1r (**Fig. 8**). These data, in combination with previous mutants of the K1 site of BBK32-C and FbpA-C (20, 31), demonstrate that residues in dynamic regions of borrelial C1r inhibitors contribute significantly to their inhibitory functions.

We speculate that dynamics of the K1 and/or K2 sites may contribute to the selectivity of borrelial C1r inhibitors for specific conformations of the protease (*i.e.*, zymogen vs. active) as has been reported for FbpB (20). However, additional study is required to fully understand this relationship that includes consideration of protein dynamics of the proteases themselves, particularly within the context of the larger C1 complex (*i.e.*, C1qC1r_2_C1s_2_). While we have shown here that FbpC does not inhibit a closely related complement pathway (*i.e.*, LP; **Fig. S3**), it is possible that the sequence diversity across BBK32/Fbp proteins may generate proteins that are capable of inhibiting more distantly related host serine proteases. Furthermore, complement proteins are highly conserved across vertebrates, including for C1r. However, we have previously noted that differences within vertebrate C1r-SP are concentrated in loop structures such as the B-loop. We hypothesize that borrelial pathogens, which naturally encounter disparate vertebrate reservoirs, may have evolved borrelial C1r inhibitors that are evolutionarily optimized for particular C1r sequences. While this possibility remains to be formally tested, evidence for this type of molecular host adaptation has been recently demonstrated for a different borrelial anti-complement lipoprotein that can bind human factor H, notably CspZ (69–71). Future work in this area should consider how protein dynamics contributes to host adaption for vector-borne pathogens like Lyme disease and relapsing fever spirochetes.

In summary, we characterized the structure and function of the C1r-binding domain of *B. hermsii* FbpC and carried out a detailed MD simulation study for the family of borrelial C1r inhibitors. This investigation reveals an unexpected plasticity in the structures of these proteins and uncovers the existence of dynamic motions within key functional sites. These observations improve our understanding of how outer surface bacterial proteins modulate the host immune response and provide novel insight into how protein dynamics may influence their functions.

## Experimental Procedures

### Plasmid Constructs

The following constructs for *B. hermsii* HS1 FbpC (GenBank #: ADN26265.2) were subcloned into pT7HMT (72). *E. coli* codon-optimized DNA fragments for the C-terminal complement inhibitory domain of FbpC and site-directed mutants (FbpC-C, FbpC-C R248A, and FbpC-C H252A) corresponding to residues 212-374 were obtained from IDT Technologies gBlock Gene Fragment Service. Residue numbering is based on UniProt numbering (G9BXS5). Each FbpC-C DNA fragment was subcloned into pT7HMT as previously described (20, 30–32).

The *B. burgdorferi* expression construct for *B. hermsii fbpC* was assembled using the NEBuilderHiFI DNA Assembly Cloning Kit as described (20, 32). Briefly, the *fbpC* gene was PCR amplified using primers BhFbpCUSF and BhFbpCDSR (**Supplementary Table 1**) from *B. hermsii* strain HS1 genomic DNA using parameters as previously indicated (20, 32). The *bbk32* promoter was amplified using primers K32PromoterUSF and K32PBhFbpCFusR **(Supplementary Table 1)**. Following assembly and confirmation by dideoxy sequencing (not shown), the entire construct was amplified and cloned into pBBE22*luc* as previously outlined (20, 32). Transcriptional linkage of the *B. hermsii fbpC* gene with the *bbk32* promoter from *B. burgdorferi* ensured transcription of *fbpC* in B314. The resulting plasmid, which encodes the P*_bbk32_*-*fbpC* construct, was designated pAP2.

### Transformation of B. burgdorferi

Transformation of strain *B. burgdorferi* B314 with the plasmid construct pAP2 was performed as previously described (31, 73). The presence of plasmid pAP2 was selected in complete BSK-II media using kanamycin at a final concentration of 300 μg/ml.

### Immunoblots

Western immunoblots were conducted to detect FbpC and BBK32 from constructs produced in *B. burgdorferi* strain B314. Protein lysates were resolved by 12.5% SDS-PAGE and transferred to polyvinylidene difluoride membranes. Membranes were incubated with either a polyclonal mouse antibody to *B. hermsii* FbpC (diluted 1:5,000), a monoclonal antibody to BBK32 (diluted at 1:20,000) or *B. burgdorferi* strain B31 FlaB (diluted at 1:10,000; Affinity BioReagents), washed, and then incubated with a 1:10,000 dilution of goat anti-mouse IgG-HRP (Thermo Fisher Scientific). The blots were processed to visualize the immune complexes as previously described (31).

### Proteinase K Accessibility Assays

Surface expression of *B. hermsii* FbpC in strain B314 was determined using proteinase K accessibility assays as described (20) with a polyclonal antibody against FbpC-C. The specificity of the polyclonal FbpC antibody was tested against strains B314-pBBE22luc (vector control) and B314-pCD100 (produces BBK32). A monoclonal antibody to BBK32 was used to confirm that *bbk32* was not being expressed and to test whether this BBK32-specific monoclonal crossreacted with FbpC when produced in B314-pAP2.

### Protein production and purification

Recombinant FbpC-C, FbpC-C R248A, and FbpC-C H252A proteins were expressed and purified as in (20). Briefly, after elution of the Ni column using the Elution buffer (20 mM Tris (pH 8.0), 500 mM NaCl, 500 mM Imidazole), the proteins were exchanged into 20 mM Tris (pH 8.0), 500 mM NaCl, 10 mM Imidazole using a Desalting 26/10 column (GE Healthcare). The His-tag was then removed by incubation with the tobacco etch virus (TEV) protease and 5 mM β-mercaptoethanol overnight at room temperature. The TEV digested proteins were separated from the cleaved His-tag product on an AKTA Pure 25L FPLC using a 5 mL HisTrap-FF with the captured flowthrough further purified using a HiLoad Superdex 75 PG gel filtration column (GE Healthcare). A monodisperse peak was obtained and assessed for purity using SDS-PAGE gel analysis, then pooled, concentrated, and exchanged into HBS buffer (10 mM HEPES [pH 7.3], 140 mM NaCl). Purified full length zymogen and active forms of C1r were obtained from Complement Technology, Inc. (Tyler, TX).

### Generation of Polyclonal Antibodies against B. hermsii FbpC

Polyclonal antibodies reactive with *B. hermsii* FbpC were generated by immunizing female C57BL/6 mice intradermally with 25 µg of purified FbpC-C in an equal volume of PBS and TiterMax^®^ Gold Adjuvant (Sigma Aldrich) as outlined (74). Two weeks after the initial immunization, mice were boosted with 25 µg of FbpC-C. Two weeks after boosting, mice were euthanized and blood was immediately isolated by exsanguination, allowed to clot, and serum removed after low-speed centrifugation. Reactivity and specificity of the serum sample to *B. hermsii* FbpC was evaluated by Western Blot with B314-pAP2 relative to the vector only control B314-pBBE22*luc*. All animal work was reviewed and approved by the Texas A&M University Institutional Animal Care and Use Committee (protocol number 2019-0422).

### Crystallization, structure determination, refinement, and analysis

FbpC-C_212-374_ was concentrated to 8 mg/mL in 10 mM Tris (pH 8.0), 50 mM NaCl buffer. Crystals of FbpC-C were obtained by vapor diffusion of sitting drops at 25°C. Crystals grew in a condition containing 0.2M MgCl_2_, 0.1M Bis-Tris (pH 5.5), 25% PEG 3,350, and generally appeared in 3 days. Crystals were harvested and cryoprotected by supplementing the crystallization buffer with 10% glycerol.

Monochromatic X-ray diffraction data were collected at 1.0Å wavelength using beamline BM-22 of the Advanced Photon Source (Argonne National Laboratory). Diffraction data were integrated, scaled, and reduced using the HKL2000 software suite (75) and assessed for data quality (76) (**Table 1**). FbpC-C crystals grew in space group P 2_1_ and contained a single copy within the asymmetric unit. Initial phases for FbpC-C were obtained by molecular replacement using the predicted FbpC-C structure from AlphaFold2 (**Fig. S1)** (35). Following an initial round of refinement, the model for FbpC-C was completed by a combination of automated chain tracing using PHENIX.AUTOBUILD (75, 77, 78) and manual building using COOT (79). The final model was completed upon iterative cycles of manual rebuilding and refinement using PHENIX.REFINE (75, 77, 78). Residues 247-254 and 301-304 were not modeled in the final refined structure due to poor electron density. Refined coordinates and structure factors have been deposited in the Protein Data Bank, Research Collaboratory for Structural Bioinformatics, Rutgers University (www.rcsb.org/) under the PDB ID 8EC3. A description of crystal cell constants, diffraction data quality, and properties of the final refined model can be found in **Table 1**. Representations of the protein structures and electron density maps were generated by PyMol (www.pymol.org/).

### AlphaFold2 Modelling

AlphaFold2 was used to predict the protein structure for *B. hermsii* FbpC_212-374_ (35) using an AlphaFold2 Colab implementation (80) and subjected to Amber relaxation. The corresponding pLDDT values were used to evaluate residue-level confidence in the predicted structure (35).

### Surface plasmon resonance

Surface plasmon resonance (SPR) was performed on a Biacore T200 instrument at 25°C using a flowrate of 30 μL/min. HBS-T running buffer was used (10 mM HEPES (pH 7.3), 140 mM NaCl, and 0.005% Tween 20) supplemented with 5 mM CaCl_2_. FbpC-C was immobilized on a CMD200 sensor chip (Xantec Bioanalytics) via standard amine coupling as before (20, 30–32). The immobilization density for FbpC-C was 1,138.2 RU, FbpC-C R248A was 975.2 RU, and FbpC-C H252A was 1,082 RU. Zymogen or active C1r binding data was obtained using a single cycle injection strategy (20) whereby a fivefold dilution series of either C1r zymogen or active C1r ranging from 0-100 nM was injected over the immobilized FbpC-C biosensor. Data was obtained in triplicate with each cycle having association times of 5 min, and a final dissociation time of 1 hour. Surfaces were regenerated to baseline with two 1 min injections with regeneration buffer (0.1 M Glycine [pH 2.2], 2.5 M NaCl). Equilibrium dissociation constants (*K*_D_) and kinetic rates were obtained from the resulting kinetic fits obtained using BiacoreT200 Evaluation Software (GE Healthcare).

### ELISA-based complement inhibition assay

ELISA-based assays that utilized IgM as a specific activator of the classical pathway or mannan as an activator for the lectin pathway as described in (20, 30–32, 81) were used to assess complement inhibition by FbpC-C or the site-directed FbpC-C mutants. Briefly, FbpC-C, FbpC-C R248A, or FbpC-C H252A were serially diluted (two-fold dilution range: 1,000 nM to 0.5 nM) and incubated with 2% normal human serum (NHS) (Innovative Research) for CP-specific ELISAs. ELISAs specific for the lectin pathway contained single 1 μM concentrations of FbpC-C or BBK32-C incubated with 2% C1q-depleted NHS (Complement Technologies). Serum incubations were performed in complement ELISA buffer (10 mM HEPES pH 7.3, 140 mM NaCl, 2 mM CaCl_2_, 0.5 mM MgCl_2_, and 0.1% gelatin). A primary mouse antibody α-C4c (Quidel) was diluted to 1:10,000 followed by a secondary goat α-mouse antibody conjugated to horseradish peroxidase (Thermo Scientific) at a 1:3,000 dilution to determine complement activation by measurement of C4b deposition. Data were obtained in at least triplicate and normalized to a well containing serum only (100% complement activation) and background subtracted using a negative control with buffer containing no serum. Plates were washed three times between each step with tris-buffered saline (50 nM Tris pH 8.0, 150 nM NaCl, and 0.05% Tween 20). An EnSight Multimode Plate Reader (PerkinElmer) was used to read the absorbance values at 450 nm for the reaction. Half-maximal inhibitory concentration values (IC_50_) were obtained using the non-linear variable slope regression fit from GraphPad Prism version 9.5.

### Serum complement sensitivity assay

Assays were performed as outlined previously (20, 31, 32). Briefly, *B. burgdorferi* strain B314 producing *B. hermsii* FbpC (pAP2), as well as the vector-only control B314 pBBE22*luc*, were grown at 32°C in 1% CO_2_ in complete BSK-II media with kanamycin at 300 μg/ml to early-to mid-log phase. The cells were then diluted in complete BSK-II media to a final cell density of 1x10^6^ cells/mL, then incubated for 1.5 hours in 15% normal human serum (Complement Technology) or equivalent heat inactivated serum as a positive control for survival. Cell survival was assessed by dark field microscopy based on cell motility and membrane damage or lysis.

### Molecular Dynamics Simulations

All atom MD simulations for the complement inhibitory domains of BBK32 (BBK32-C) from *B. burgdorferi* and orthologous proteins FbpA (FbpA-C), FbpB (FbpB-C) from *B. miyamotoi* and FbpC from *B. hermsii* (FbpC-C) were carried out with coordinates from their solved crystal structures (PDB IDs: 6N1L, 7RPR, 7RPS and 8EC3, respectively). Residues not visualized in the final crystal structures due to poor electron density were first modelled with MODELLER v9.1 (82). OPLS-AA force fields (83) were used with explicit TIP3P water model for all simulations. All atom models for the respective proteins were constructed with hydrogens and all simulations were performed with Gromacs v 2021.3 (84). A total of 24,698, 24,283, 23,751 and 33,073 water molecules were used to solvate BBK32-C, FbpA-C, FbpB-C and FbpC-C systems respectively. An application of periodic boundary conditions of 10 Å from the edge of the cubic box of dimensions 91.3, 90.8, 90.2 and 100.5 Å, respectively. Charge neutralization of the simulation systems for BBK32-C, FbpA-C, FbpB-C and FbpC-C were undertaken by replacing water molecules with 5 Cl-, 3 Cl-, 1 Cl- and 2 Cl-ions, respectively. The final systems to be simulated were thus comprised of 76,440 atoms for BBK32-C, 75,355 atoms for FbpA-C, 73,926 atoms for FbpB-C, and 101,858 atoms for FbpC-C. Bond lengths for each of these systems were constrained by LINCS algorithm (85) and all long-range electrostatic interactions were determined using the smooth particle mesh Ewald (PME) method (86). Energy minimization was performed with a steepest descent algorithm until convergence (∼1,000 steps) with a maximum number of steps set to 50,000. All simulations were performed at 300K. Temperature equilibration was conducted by the isochoric-isothermal NVT ensemble (constant number of particles, volume, and temperature) with a Berendsen thermostat for 100 ps (87). The system was then subjected to pressure equilibration in the NPT ensemble (constant number of particles, pressure, and temperature) for 100 ps using the Parrinello–Rahman protocol (88), maintaining a pressure of 1 bar. Coordinates were saved at an interval of 10 ps, and backbone RMSD, RMSF, radius of gyration and trajectory analyses were performed with GROMACS programs ‘gmx rms’, ‘gmx rmsf’,’gmx_gyrate’ and ‘gmx trjconv’ (89, 90). All simulations were performed for a period of 1 µs and attained convergence based on Cα RMSD and visual inspection. Structural snapshots were extracted from the trajectories at an interval of 100 ps leading to 10,000 snapshots for all simulations. All simulations were performed on a local workstation with CUDA acceleration v11.2 powered by an NVIDIA Quadro GPU with 2560 CUDA compute cores resulting in an average output simulation trajectory of ∼ 50 ns/day. DCCM and normal mode analyses (all atom and Cα) were performed with Bio3D package (91). 2D free energy surface generation with reaction coordinates of side chain and main chain distances between the atoms of K1 and K2 site (defined as K1:R248 and K2:K327 for BBK32-C, K1:R264 and K2:K343 for FbpA-C, K1:R309 and K2:N402 for FbpB-C.

Two sites were measured for FbpC-C; K1:R248 and K2:K345 and K1:H252 and K2:K345. Coordinates were separated into 100 bins followed by discrete probability distribution calculations according to the equation below, where P*_i_* is the probability in a particular bin and P*_max_* the maximal probability, R is the universal gas constant while temperature (T) is measured in K (298 K) at which all the simulations were performed (82).

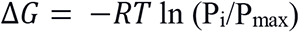

### DCCM Analysis

Formal definition of dynamic cross correlation matrices is given by the equation below where r_i_ and, r_j_ are the spatial backbone atom positions of the respective i^th^ and j^th^ amino acids (83):

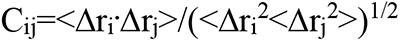

A time scale is associated with each C_ij_ element, and this time scale corresponds to a subset of contiguous snapshot structures taken from the temporal series of snap-shot structures extracted from the MD trajectory. This timescale (subset of structures) determines the time interval over which the C_ij_ elements are calculated. Positive C_ij_ values result from backbone atom motions between residues i and j that are in the same direction along a given spatial coordinate; while negative C_ij_ values result from backbone atom motions between residues i and j that are opposite in direction along a given spatial coordinate. To normalize such positional fluctuation, all the MD snapshot frames extracted for calculations were superposed to a common frame of reference (*i.e.*, the crystal structure for each protein).

## Data Availability

The datasets presented in this study can be found in online repositories. The names of the repository/repositories and accession number(s) can be found below: http://www.wwpdb.org/, 8EC3. All other data are contained within the manuscript.

## Supporting Information

This article contains supporting information.

## Author Contributions

S.R. and C.E.B. are equal contributors. J.T.S. and B.L.G. are co-corresponding authors. S.R., C.E.B., A.D.P., A.S., J.T.S and B.L.G. designed research; S.R., C.E.B., A.D.P., and A.S., conducted research; S.R., C.E.B., A.D.P., A.S., J.T.S and B.L.G analyzed data; S.R., C.E.B., A.D.P., J.T.S and B.L.G wrote the paper.

## Funding and additional information

Support was provided by Public Health Service Grant R01-AI146930 from the National Institute of Allergy and Infectious Diseases (B.L.G and J.T.S.). The content is solely the responsibility of the authors and does not necessarily represent the official views of the National Institutes of Health. X-ray diffraction data were collected at Southeast Regional Collaborative Access Team 22-BM beamline at the Advanced Photon Source, Argonne National Laboratory. Supporting institutions may be found at www.ser-cat.org/members.html.

