## Supplemental Information for "“Conformational dynamics of C1r inhibitor proteins from Lyme disease and relapsing fever spirochetes”"

Running Title: BBK32/Fbps MD simulations

Sourav Roy[1†], Charles E. Booth Jr.[1†], Alexandra D. Powell-Pierce[2], Anna M. Schulz[1], Jon T. Skare[2*], and Brandon L. Garcia[1*]

[1]Department of Microbiology and Immunology, Brody School of Medicine, East Carolina University, Greenville, North Carolina, United States of America

[2]Department of Microbial Pathogenesis and Immunology, College of Medicine, Texas A&M University, Bryan, TX, United States of America

[†]Equal contribution

**This file contains Figures S1-S5 and Tables S1-S2**

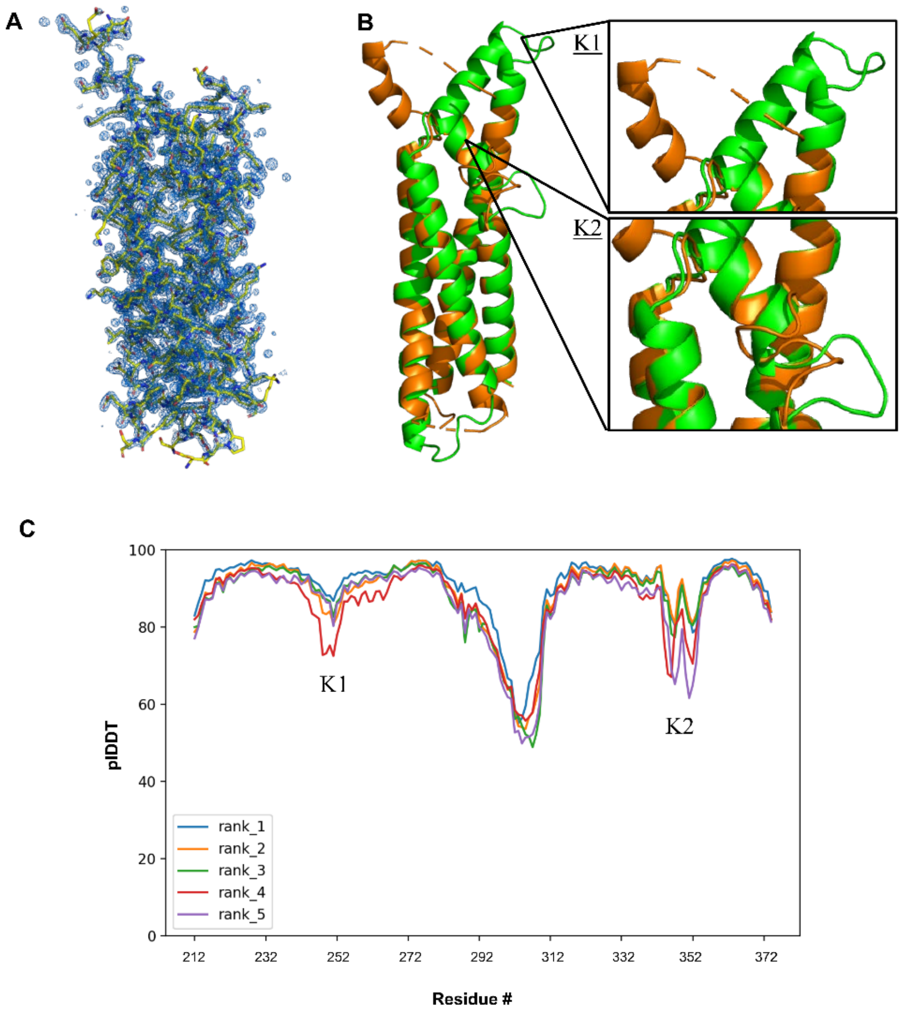


**Supplemental Figure 1. *B. hermsii* FbpC-C electron density maps and analysis of the AlphaFold2 model. A)** The crystal structure of *B. hermsii* FbpC-C$_{212\text{-}374}$ depicted as yellow sticks with the corresponding *2Fo-Fc* electron density map contoured to 1.5 σ. **(B)** A structural alignment of the crystal structure of FbpC-C (PDB ID: 8EC3) (orange) and the AlphaFold2 model used for molecular replacement (green). The AlphaFold2 model differs from the crystal structure in the arrangement of the second alpha helix. The K1 and K2 sites are shown as inlets to the right. **(C)** FbpC-C was folded using AlphaFold2 Colab (35, 80) producing five models. The residue-level confidence metric, plDDT, is plotted and shows that lower confidence predictions are made for residues in the K1 and K2 sites of FbpC-C.

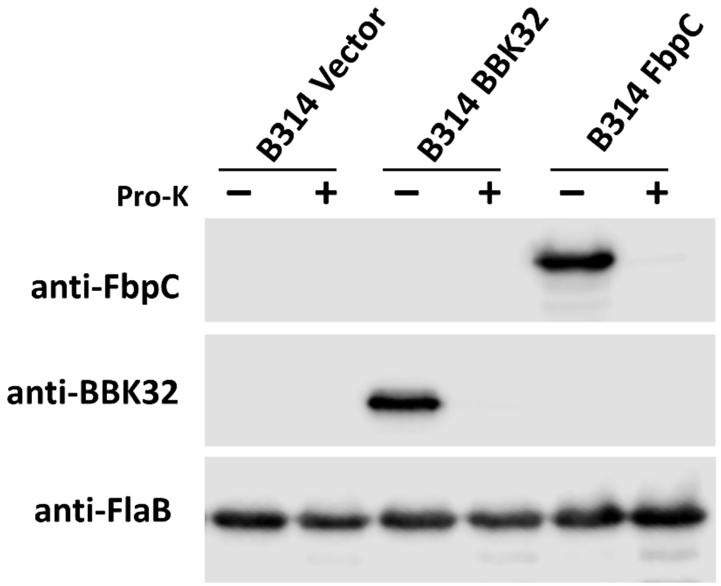


**Supplemental Figure 2. Surface expression of *B. hermsii* FbpC in *B. burgdorferi* B314.** Proteinase K accessibility assays were used to determine the surface localization of FbpC in *B. burgdorferi* strain B314 transformed with plasmid pAP2 (B314 FbpC) using a polyclonal antibody against FbpC. FbpC is not present nor does it cross react with any *B. burgdorferi* protein in B314 pBBE22*luc* (vector control) or B314 pCD100 (BBK32). A monoclonal antibody to BBK32 did not recognize any antigen in B314 FbpC strain, indicating that *B. hermsii* FbpC does not contain the monoclonal antibody-specific BBK32 epitope. Protein expression is normalized to a flagellar loading control, FlaB.

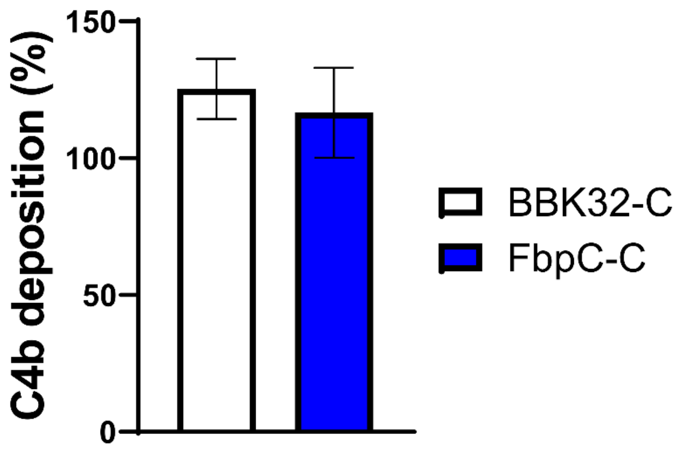


**Supplemental Figure 3. Lectin pathway-specific inhibitory complement ELISA.** Mannan was immobilized on an ELISA plate and C1q-depleted normal human serum incubated with 1 µM FbpC-C or BBK32-C. Neither FbpC-C nor BBK32-C inhibited the lectin pathway as indicated by the deposition of C4b. Data was obtained in triplicate and shown as the average of the replicates with the standard error.

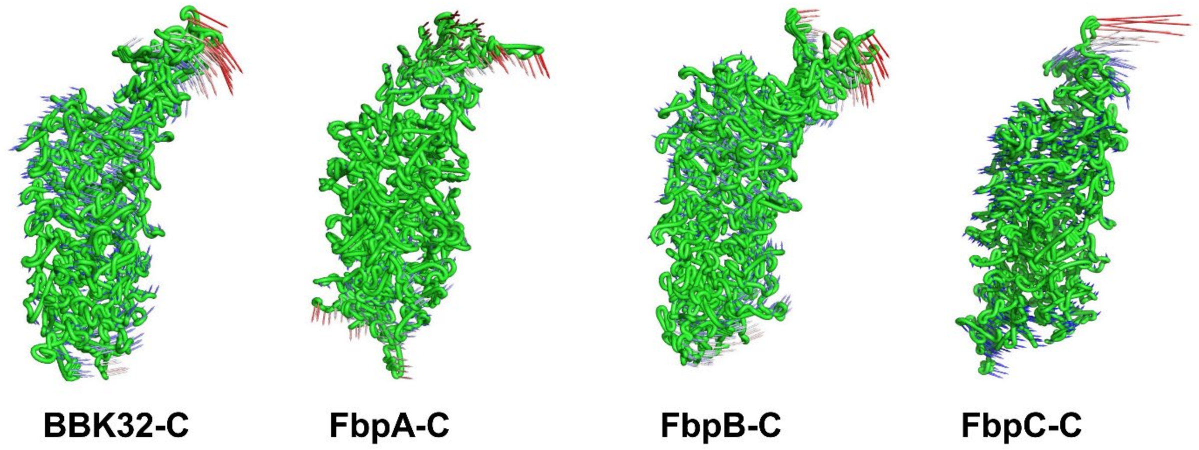


**Supplemental Figure 4. Normal mode analysis for BBK32-C, FbpA-C, FbpB-C, and FbpC-C.** All-atom Normal modes for BBK32-C, FbpA-C, FbpB-C and FbpC-C demonstrating large amplitude vibrations marked in red around K1 site with smaller amplitude anti-correlated motions around K2 site marked in blue.

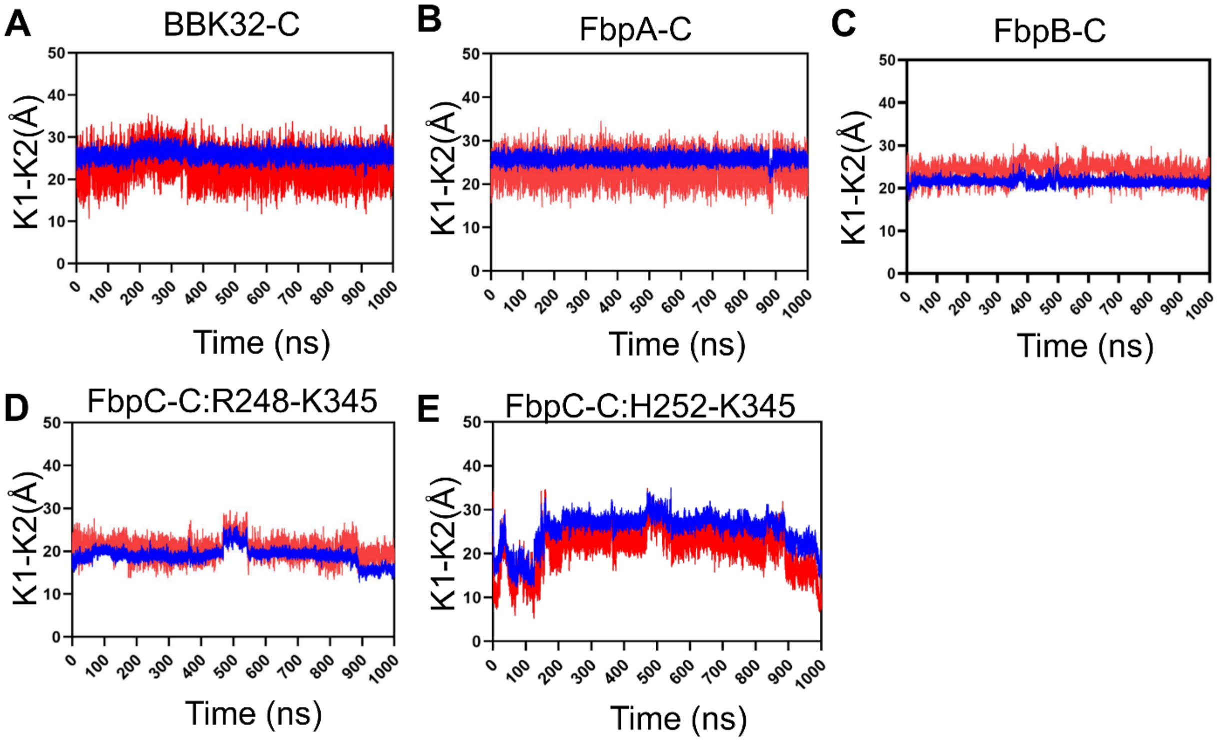


**Supplementary Figure 5. Distance fluctuations between the K1 and K2 sites.** Main chain (red) and side chain (blue) distances between K1 and K2 sites are tracked over the time course of a representative simulation. The residues used for these measurements are defined as **(A)** BBK32-C (K1: R248, K2: K327), **(B)** FbpA-C (K1: R264, K2: K343), and **(C)** FbpB-C (K1: R309, K2: N402). Two measurements were tracked for FbpC-C: **(D)** (K1: R248, K2: K345) and **(E)** (K1: H252, K2: K345).

| **Supplementary Table 1. Oligonucleotides used for the present study.** | | | |
|---|---|---|---|
| **Oligonucleotides** | **Sequence** | **Description** | **Reference** |
| K32PromoterUSF | GAGGTACCCGGGGATCCGTACTTTGTTCACCCTCTTG | Oligonucleotide pair used to amplify the *bbk32* promoter for assembly with the *fbpC* pBBE22*luc* to produce pAP2 | (32) |
| K32PBhFbpCFusR | CATTGTTTTTCAATTGCATGCTTTCTCTCCTTTAAAGTTAATAC | | This study |
| BhFbpCUSF | GTATTAACTTTAAAGGAGAGAAAGCATGCAATTGAAAAAACAATG | Oligonucleotide pair used to amplify *B. hermsii fbpC* for assembly with the *bbk32* promoter and pBBE22*luc* to produce pAP2 | This study |
| BhFbCDSR | GCTTGCATGCCTGCAGGTCGACTTATTTTAAGGCTTTCTTG | | This study |

| **Supplementary Table 2.** *Borreliella burgdorferi* strains used for the present study. | | |
|---|---|---|
| ***Borreliella burgdorferi* Strain** | **Description** | **Reference** |
| B314 | Serum-sensitive, non-infectious *B. burgdorferi* B31 derivative strain lacking all linear plasmids. | (37) |
| B314 pBBE22*luc* | B314 with shuttle vector encoding *bbe22* and *B. burgdorferi* codon-optimized *luc* gene under the control of a strong borrelial promoter ($P_{flaB}$-*luc*); $kan^R$. | (29) |
| B314 pCD100 | B314 with *bbk32* under control of its native promoter in pBBE22*luc*; $kan^R$. | (31) |
| B314 pAP2 | B314 with *B. hermsii fbpC* under control of the *B. burgdorferi bbk32* promoter in pBBE22*luc*; $kan^R$. | This study |
